# Directional motion of a vesicle on conical surfaces

**DOI:** 10.1101/2020.01.09.899997

**Authors:** Ge Zhenpeng

## Abstract

*The spontaneous directional motion of vesicles on both the outer and inner surfaces of a conical substrate is observed in this work. We showed that the motion is ultra-fast and the maximum velocity can be as high as 2.14 nm*/μs. *The driving force behind is attributed to the reduction of the bending energy along the conical surface, which possesses high curvature gradient*.

**Summary:** We observed and explained the spontaneous directional motion of vesicles on both the concave and convex surfaces of a cone.

## Introduction

Vesicle is an essential component in cell biology and perform a variety of functions. The directional transport of vesicles play critical roles in many physiological processes, such as endocytosis[1], exocytosis[2], axonal growth[3] and nerve growth[4] as well as the delivery of mRNA vaccines. The directional transport of vesicles is long-term believed to be ATP-dependent, which involves the cooperation of cytoskeleton and different motor proteins[5, 6].

In this work, we revealed a novel spontaneous mode of directional motion of vesicles, which purely driven by the curvature gradient. Specially, we discovered that a vesicle can spontaneously move from the highly-curved end to the flat end on the outer surface (Fig1 (a)), while moves in the reverse direction in the inner surface of a cone (Fig1 (b)).

**Fig 1.**
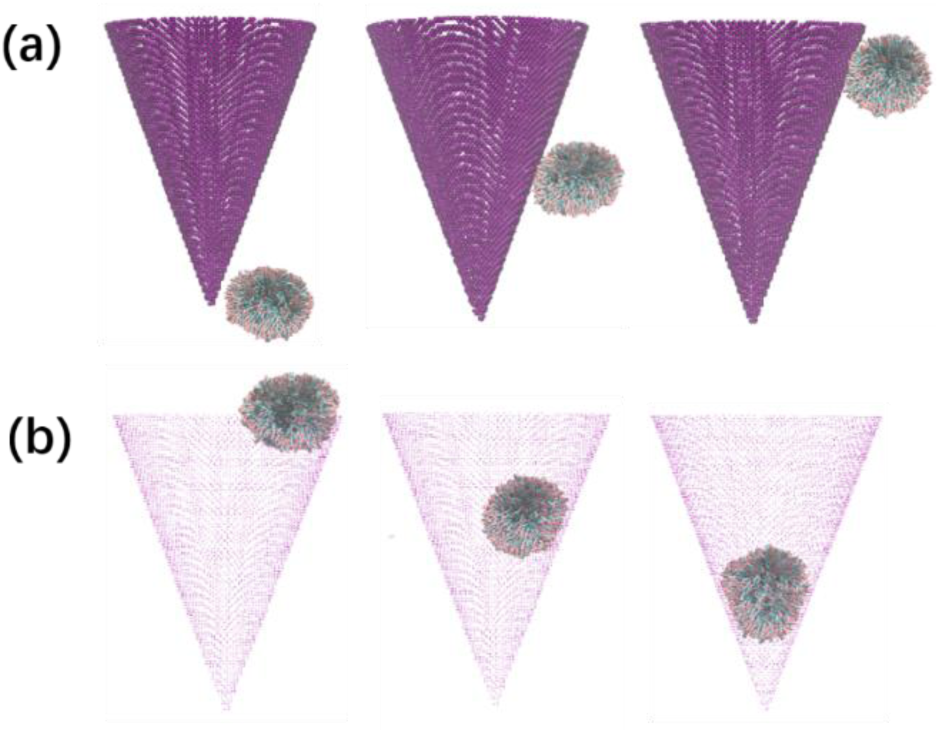
A schematic representation of the directional motion of vesicle on (a) outer surface (b) inner surface of the conical substrate. All figures are taken from simulations.

## Methods

The solvent-free force field from Cooke[7, 8] was adopted for vesicles in this work. In this model, each lipid molecular contains one head bead and two tail beads. The interactions between the beads are governed by a combination of Weeks-Chandler-Andersen potential[9], attractive potential as well as additional bonding and bending terms. The unit simulation timescale is τ ≈ 10 ns, which is estimated upon lipid self-diffusion simulations. The unit length is σ ≈ 1 nm, which is estimated by comparing the thickness of a bilayer from experiments and simulations.

The substrate is built by first constructing a cube with the face-centered cubic structure and then cutting it into a conical shell. The half-angle is set to be 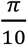 and 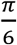, separately, to mimic different curvature gradient. The length between adjacent beads in the cube is set to be 1σ. The interaction between the substrate and vesicle is modeled as the same with that of the tail-tail interaction in vesicles, which has the following form.

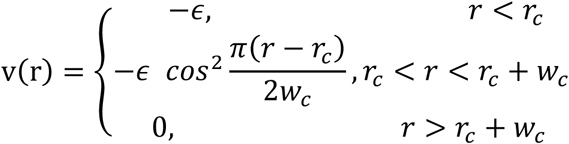

We change *ϵ* from 0.05 to 1.0 to study the effect of binding strength between the vesicle and substrate with a higher *ϵ* corresponding to stronger binding strength. The diameter of the vesicle in this study is set to be 6 nm, which ensures its stability in solution.

**Table 1.**
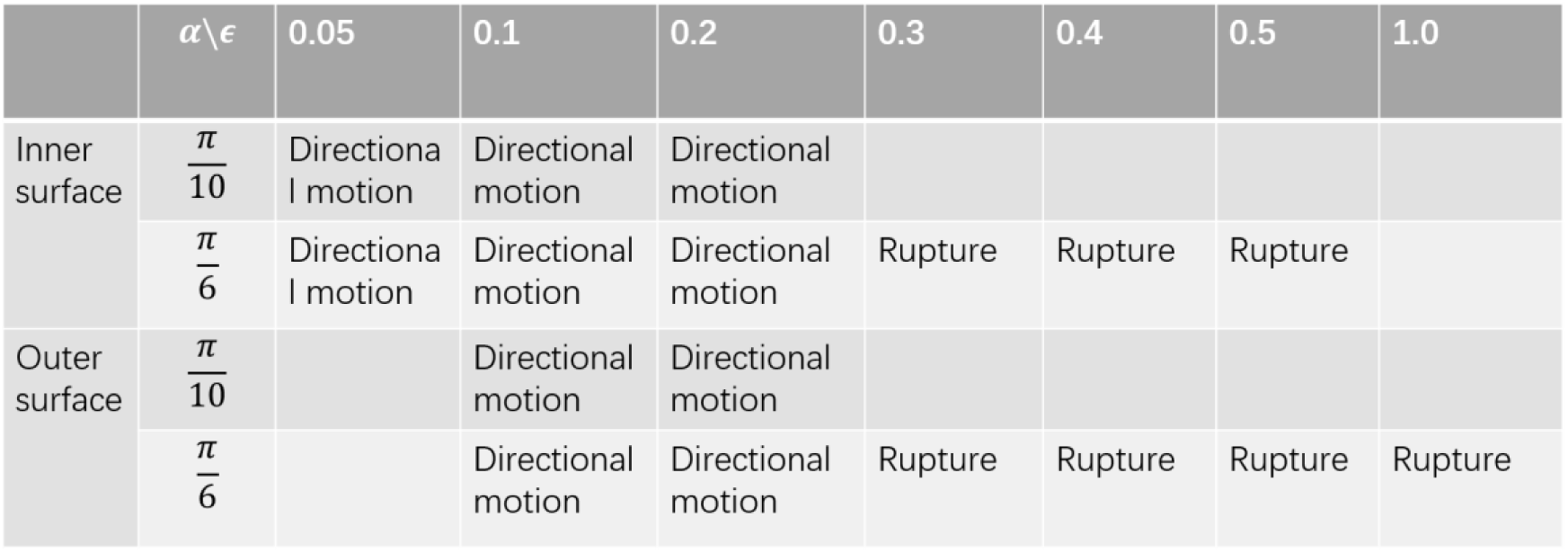
The results of the systems studied in this work.

## Results and discussions

The directional motion of vesicles is observed on both the outer and inner surface of the conical substrate. We will describe them in detail separately.

## Outer surface

As mentioned earlier, the half-angle of the conical substrate is set to be 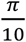 and 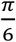, separately. For each system, we changed *ϵ* to mimic different binding strength between the vesicle and the substrate.

In the system with half-angle being 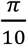, directional motion was observed when *ϵ* = 0.1 and 0.2. As shown in Fig 2, the maximum climbing height is ∼69 nm when *ϵ* is 0.1, which is much larger than that (∼47 nm) when *ϵ* is 0.2. The maximum velocity in the former system is also larger than the latter, which is 2.14 and 1.56 nm/μs, separately.

**Fig 2.**
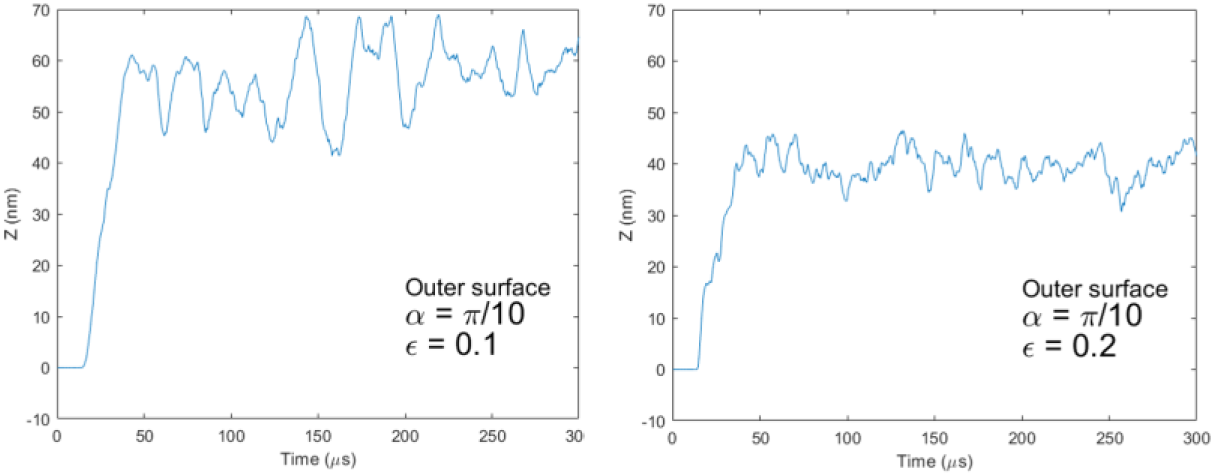
The relationship between simulation time and Z coordinate of the vesicle when the vesicle is on the outer surface and the half-angle of the cone is 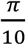.

In another system with half-angle being 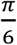 (Fig 3), we built more sub-systems with *ϵ* ranging from 0.1 to 1.0. We oberserved vesicle direntional motion when *ϵ* = 0.1 and 0.2. When *ϵ* = 0.1, the vesicle climbed to a higher position compared to the case when *ϵ* = 0.2, which is consistent with the observation in the system with half-angle being 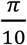. We observed the step-wise climb of the vesicle for both *ϵ* = 0.1 and 0.2.

**Fig 3.**
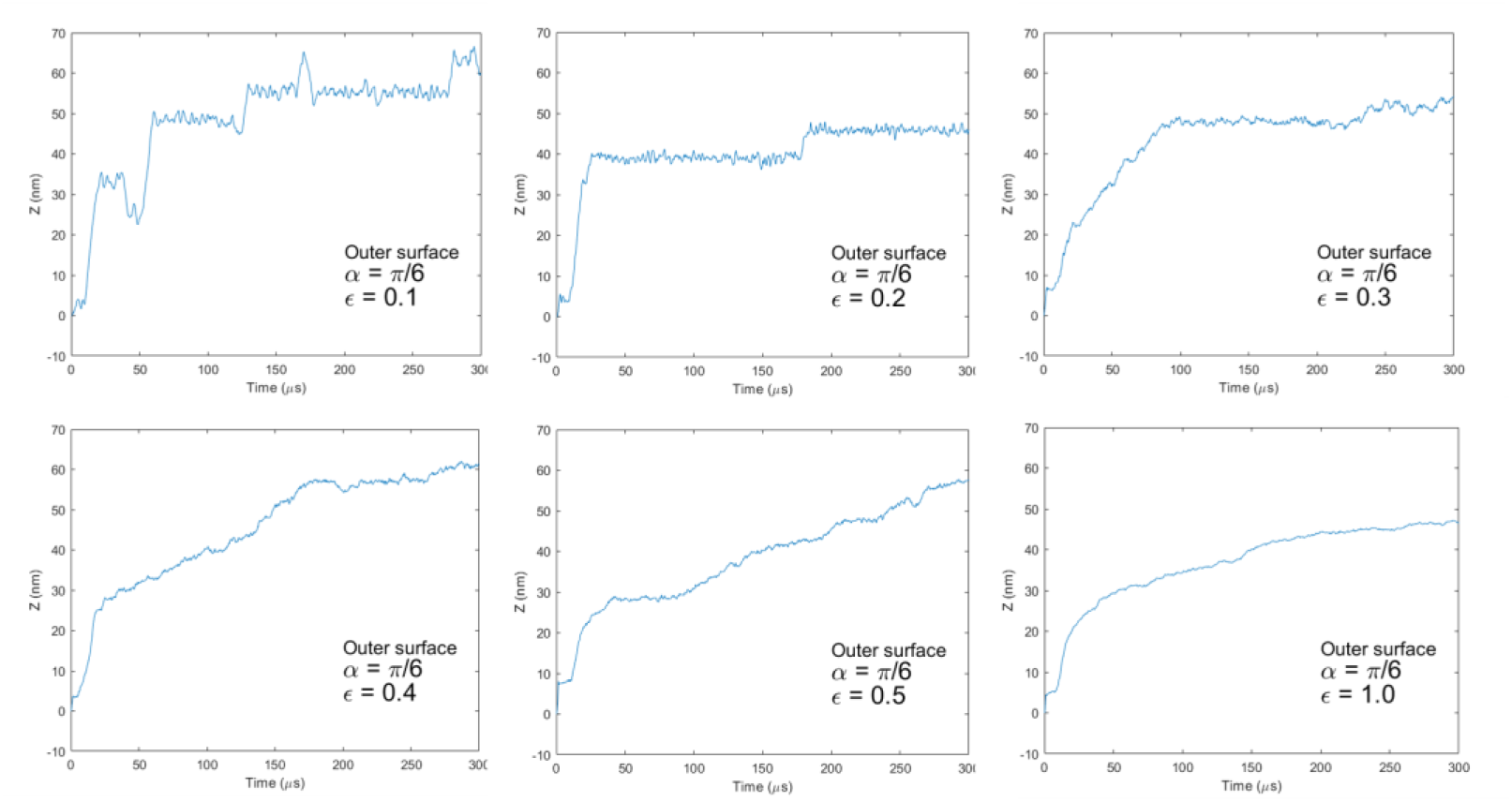
The relationship between simulation time and Z coordinate of the vesicle when the vesicle is on the outer surface and the half-angle of the cone is 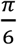.

The vesicle first took very short time to climb and then “rest” there for a relatively longer time and then climb again. This process may repeat for several times.

The vesicle ruptured and became a flat and round disc when *ϵ* = 0.3 or larger (Fig 4 (a)). What’s interesting is that ruptured disc also directionally moved towards the top flat region of the conical substrate. The maximum height of the disc when *ϵ* = and 0.5 is a little bit larger than that of *ϵ* = 0.3 and 1.0. The disc climbed faster when *ϵ* = 0.3 compared to other systems.

**Fig 4.**
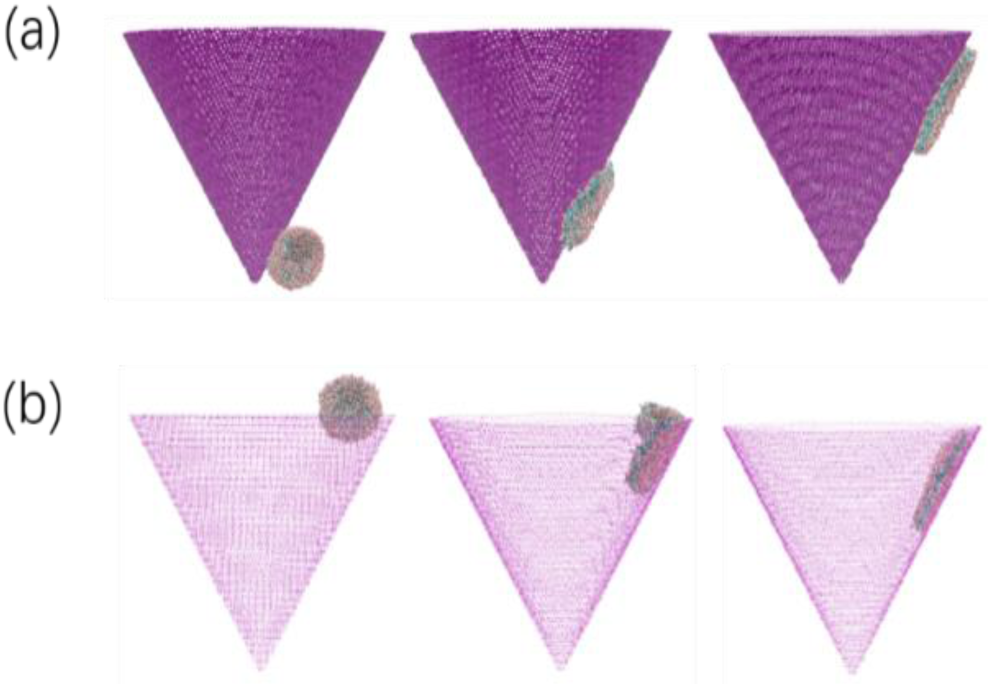
The rupture of vesicles on (a) outer surface and (b) inner surface of a conical surface.

## Inner surface

When the half angle is 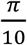, the vesicle directionally moved from the top to the bottom of the inner surface (Fig 5). The vesicle takes fewer time when *ϵ* = 0.05 compared to that when *ϵ* = 0.1 or 0.2.

**Fig 5.**
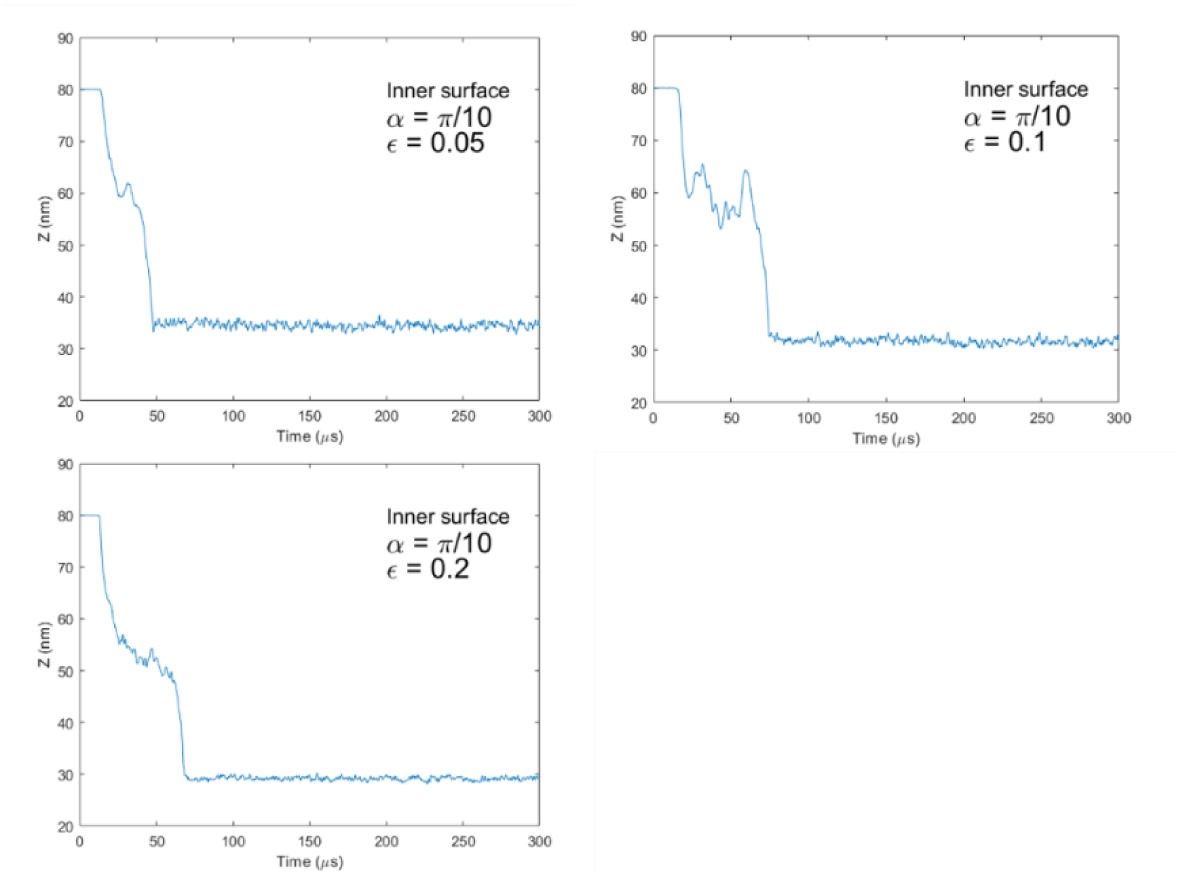
The relationship between simulation time and Z coordinate of the vesicle when the vesicle is on the inner surface and the half-angle of the cone is 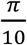.

When half angle is 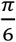, we also have more sub-systems with *ϵ* ranging from 0.05 to (Fig 6). The vesicle directionally moves on the inner surface when *ϵ* = 0.05, 0.1 and 0.2. When *ϵ* = 0.1, the vesicle takes the shortest time to reach the bottom of the substrate. When *ϵ* = 0.2, the vesicle cannot reach the bottom during our simulation time.

**Fig 6.**
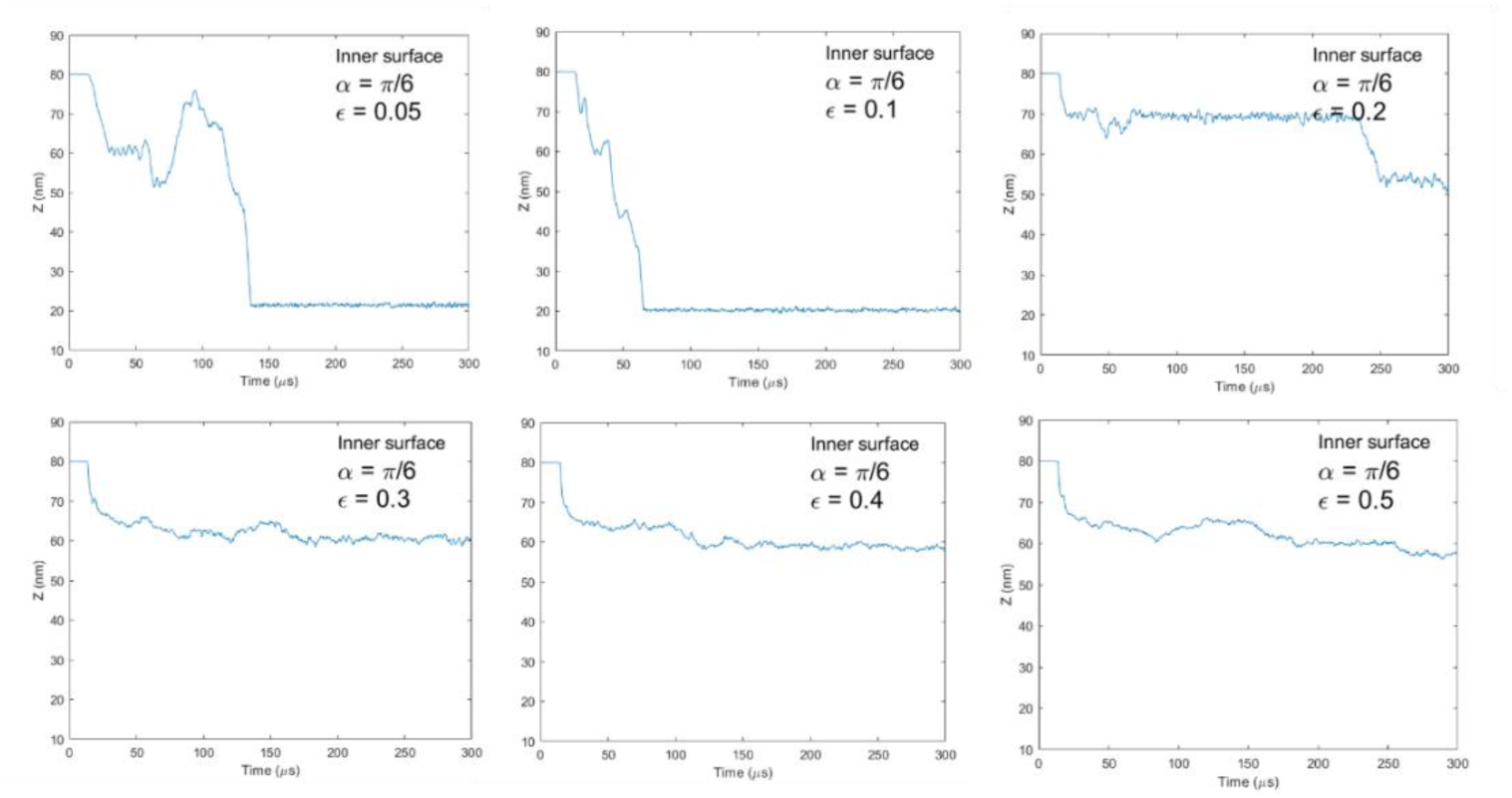
The relationship between simulation time and Z coordinate of the vesicle when the vesicle is on the inner surface and the half-angle of the cone is 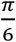.

When *ϵ* is equal to or larger than 0.3, the vesicle ruptures to a flat round disc and moves downwards with a very short distance to maximize the contact area with the substrate (Fig 4 (b)).

## Driving force

These observations can be well explained by the elastic theory developed by Helfrich[10] and Canham[11], in which the membrane is described as a ultra-thin sheet with the following energy terms:

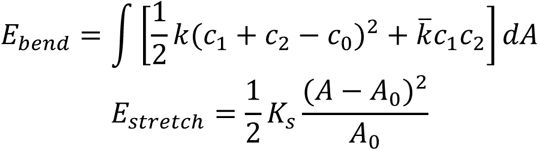

where *c*_1_ and *c*_2_ are the two principle curvatures at a given point on the membrane surface and *c*_0_ is the spontaneous curvature of the membrane; *k*, 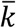 and *K*_*s*_ are the bending modulus, the saddle-splay modulus, and the stretching modulus; and *A* and *A*_0_ represent the current and the equilibrium areas of the membrane, respectively. The spontaneous curvature *c*_0_ can result from an asymmetric composition of the membrane, eg, the upper and lower leaflets of the bilayer may contain different lipid species[12]. In our simulations, *E*_*stretch*_ equals 0 because there is no osmotic pressure in our solvent free model.

The interaction between the substrate and the membrane is proportional to the contact area with the following term:

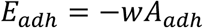

where *w* represents the binding strength between the substrate and membrane. So the total energy is

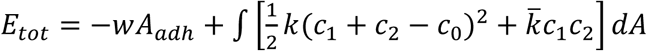

The motion of the vesicle is driven the demand to minimize *E*_*tot*_. There exists large bents at the contact point with the substrate on the vesicle when the vesicle adheres to the high-curved part on the outer surface (Fig 7), which costs high bending energy. So the vesicle moves directionally from the high-curved part to the flat part. But when the binding strength is too high, the vesicle ruptures to maximize the contact area with the substrate.

**Fig 7.**
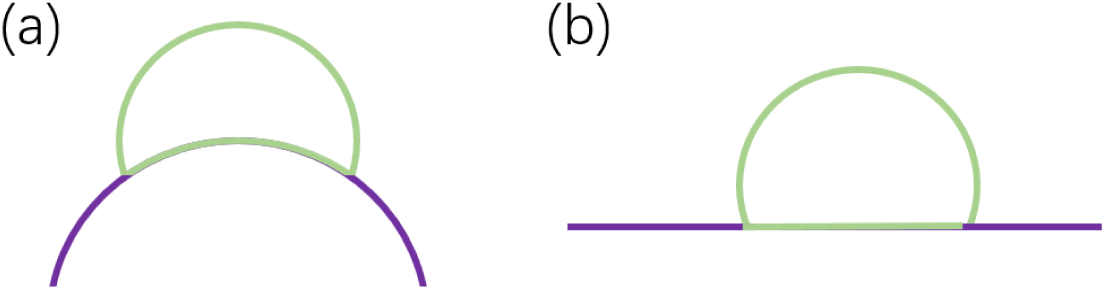
A schematic representation of a vesicle adhering to (a) the curved part and (b) flat part of outer surface of the substrate.

When the vesicle is on the inner surface of the substrate, the vesicle costs less bending energy adhering on the high-curved part (Fig 8 (a)) compared to the flat part (Fig 8 (b)). So the vesicle moves directionally from the flat part to the more curved part. Similarly, when the binding strength is too high, the vesicle ruptures to maximize the contact area with the substrate.

**Fig 8.**
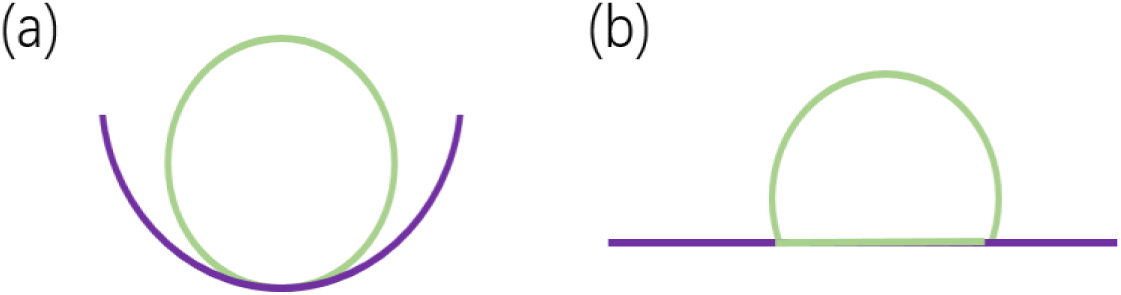
A schematic representation of a vesicle adhering to (a) the curved part and (b) flat part of inner surface of the substrate.

## Conclusions

In this wok, we observed the directional motion of a vesicle on both the outer and inner surface of a conical substrate. The motion is spontaneous and purely driven by the curvature gradient of the substrate. Besides the curvature gradient, the behavior of the vesicle is also affected by the adhesion strength between the vesicle and substrate. Upon certain threshold, the vesicle will rupture. We showed that all these phenomenon can be well explained by the elastic theory.

## Supporting information

Directional motion of a vesicle on the inner surface of a cone.

Directional motion of a vesicle on the outer surface of a cone.

---

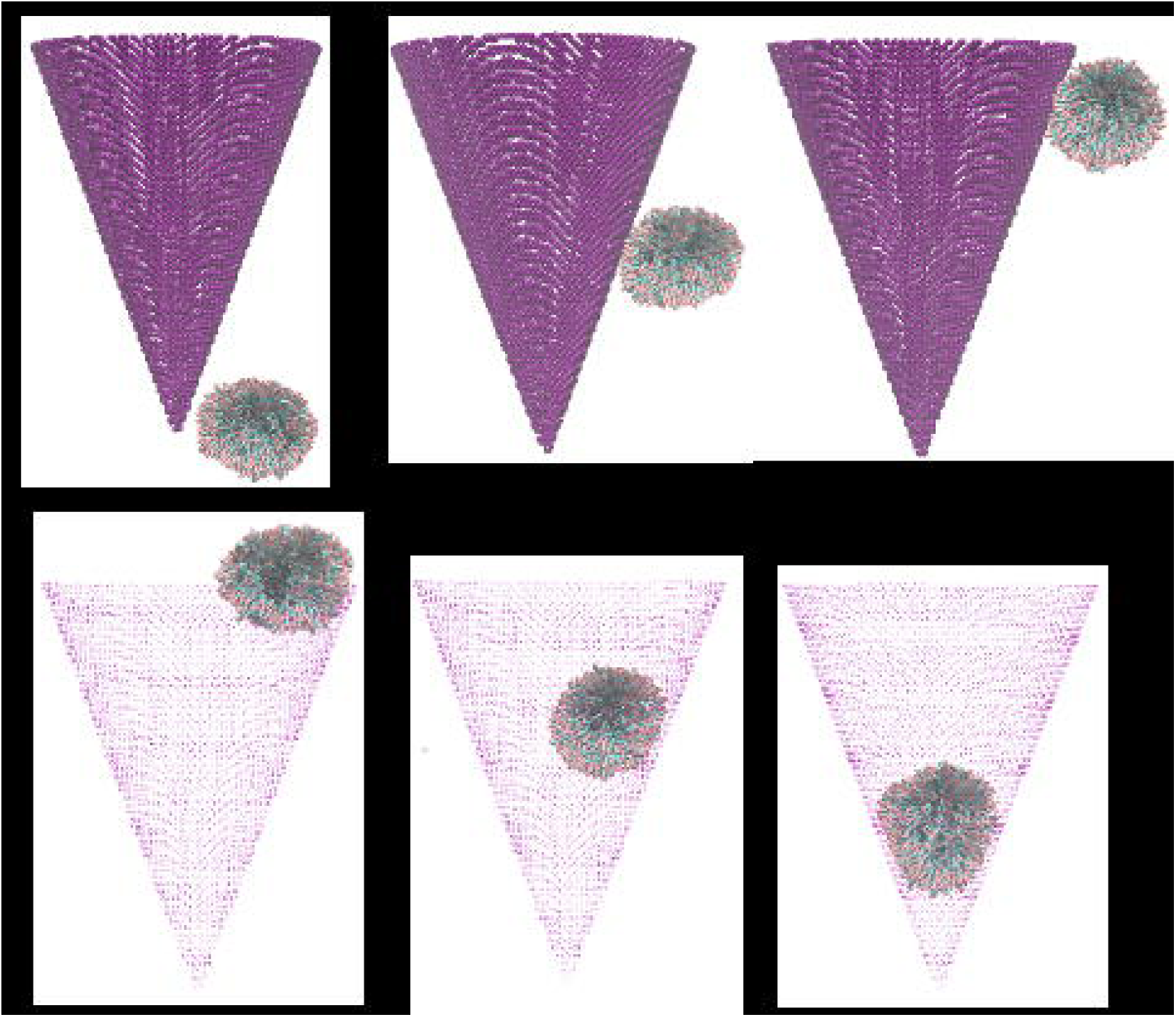

